# Inhibition of fibrosis with multi-agent therapy in pulmonary fibrosis: Results of a drug library screening

**DOI:** 10.1101/2020.06.29.178061

**Authors:** Cassandra Batzlaff Braun, Megan Girtman, Paige Jenson, Michael H Bourne, JuneMee Chae, Theodore Kottom, Andrew Limper

**Author notes:** <u>Corresponding author and reprints:</u> Dr. Andrew Limper, 8-24 Stabile Building, Mayo Clinic, Rochester, MN 55905., and.

## Abstract

**Aims:** Successful management of IPF will likely require multi-drug therapy as its pathogenesis is thought to be both driven by both pro-inflammatory and pro-fibrotic pathways. We hypothesized that the available anti-fibrotic agents, pirfenidone and nintedanib, may exhibit synergy in suppressing lung fibroblast extracellular matrix protein generation when administered in combination with other orally active agents.

**Materials and Methods:** A fibroblastic cell line (AKR-2B) was stimulated with TGF-β1 and used to screen a library of over 1500 FDA approved drugs. Extracellular matrix protein generation was assessed via fibronectin ELISA assay and maintenance of cell viability confirmed with XTT assay.

**Results:** The screening revealed sixty-two drugs from the repurposed drug-screening library that were shown to significantly suppress fibronectin expression and not result in cell death. Specifically drugs within the category of NSAIDs, steroids, azole antifungal agents, and antipyrine were associated with significant suppression of fibronectin on ELISA analysis. Surprisingly, we observed anti-fibrotic activity across a number of the azole antifungal compounds. We next assessed whether combination of azoles would exhibit synergy when combined with current anti-fibrotic therapies in the stimulated fibroblasts. As proof of concept, we demonstrated *in vitro* synergy between oxiconazole and nintedanib in suppressing fibroblast generation of extracellular matrix fibronectin.

**Conclusions:** These results suggest an approach to identify potential combinations of therapy that may improve patient outcomes by reducing cost and potential toxicities during treatment.

## Introduction

Idiopathic pulmonary fibrosis (IPF) is a chronic and progressive interstitial lung disease with limited treatment options. The prognosis for patients with this diagnosis remains quite poor with an average life expectancy of 50% after 3 to 5 years from the time of diagnosis ^1^. The pathogenesis of this disease process is not fully understood, but is thought to involve both pro-inflammatory and pro-fibrotic mechanistic pathways. On pathologic evaluation, the lungs in IPF are characterized by heterogeneous distribution of normal regions alongside areas of dense fibrosis and fibroblastic foci. This distortion of architecture results in lung remodeling with frequent honeycombing, predominantly in a subpleural or paraseptal distribution ^2^. Current mechanistic concepts indicate that repeated micro-injuries of the alveolar epithelium result in fibroblast and myofibroblast proliferation and apoptosis, driven by profibrotic growth factors including TGF-β1. These processes ultimately lead to the excessive deposition of extracellular matrix components characteristic of this disease ^3^.

The approval of two anti-fibrotic agents, pirfenidone and nintedanib, by the US Food and Drug Administration for IPF has initiated an exciting new era in the treatment of this disease. However, the clinical impact on mortality and disease progression in real world settings remains uncertain and these therapies are costly and have associated side effects. In clinical trials, both of these agents were shown to result in a statistically slower decline in forced vital capacity (FVC) over a year for those IPF patients with mild to moderate impairment in lung function ^4–7^. However, while pirfenidone was initially shown to have some improvement in all-cause mortality for the duration of the study, its effects on long-term outcomes are still uncertain ^7^. In addition, nintedanib was found to prolong the time to acute exacerbation in patients with IPF. However, nintedanib did not demonstrate any benefit in all-cause mortality ^4^.

A number of alternative agents have already been independently evaluated for IPF in clinical trials; for example imatinib and N-Acetyl Cysteine (NAC) have been studied. However, neither imatinib, nor NAC, had any effect on preservation of lung function or survival in IPF ^6,8^. Other agents that have also been tested in clinical trials for IPF without evidence of overall benefit have included bosentan, etanercept, simtuzimab, ambrisentan, anticoagulation with warfarin, and sildenafil, among others ^9^.

Many other chronic lung diseases including COPD, asthma, tuberculosis, and lung cancer have required multi-drug therapy for effective long-term disease management. Such an approach will also likely be the case for IPF since its pathogenesis is driven by both pro-inflammatory and pro-fibrotic pathways as described ^10^. The idea of utilizing a multi-drug therapy approach introduces the potential benefit of additive or synergistic effects between treatments, which may allow for use of lower doses of each individual drug and as a result fewer adverse side effects ^11^.

In light of the limited therapeutic options for pulmonary fibrosis, we undertook a drug screening strategy utilizing a library of available FDA drugs approved for other indications ^12^, to identify compound classes that might also possess anti-fibrotic activity. We hypothesized that the novel anti-fibrotic agent, nintedanib when combined with other compounds may exert synergistic activity in suppressing fibroblast mediated extracellular matrix protein generation. We posit that such an approach might be exploited to identify new multiple agent treatment regimens that may be tested in idiopathic pulmonary fibrosis.

## Materials and Methods

### Reagents

TGF-β1 was purchased from R&D systems, Minneapolis, MN, activated by acidic conditions, and then stored at −20°C until use. Unless otherwise specified, all other general reagents were from Sigma-Aldrich (St. Louis, MO).

### Cells

The AKR-2B fibroblast line was obtained from Sigma Aldrich (St Louis, MO) and maintained in McCoy’s 5A media purchased from American Type Culture Collection (ATCC, Manassas, VA) with 10% fetal bovine serum (FBS) and 1X antibiotic/antimycotic from Gibco (Grand Island, NY) (growth media). Cells were used between passages 3 and 15.

### Drug preparation

A screening library of Food and Drug Administration (FDA) approved drugs consisting of 1524 compounds representing a wide variety drugs was obtained through the Johns Hopkins Clinical Compound Library (JHCCL), version 1.3 for analysis ^12^ and assayed at 10 μM concentrations for their ability to suppress extracellular matrix fibronectin production by cultured fibroblasts (AKR-2B Cells). For additional testing beyond the screen, drug compounds were purchased from either the Mayo Clinic Pharmacy (Rochester, MN) or Sigma-Aldrich (St. Louis, MO) and solubilized in sterile DMSO or McCoy’s 5A, 0.1% FBS, 1X antibiotic/antimycotic (starve media) to make a stock solution. From the individual stock solutions a series of dilutions were then prepared for each agent for testing.

### Cell culture and compound screening

The AKR-2B cells were plated at a concentration of 10,000 cells/well on a 96 well culture plate with growth media and grown to confluence for 24 hours. Media was removed and replaced with starve media as described above for 24 hours. The above prepared drug dilutions were then applied to the culture plate for one hour and stimulated with TGF-β1 at a concentration of 5 ng/ml and incubated at 37°C for 18 hours in the presence of the test compounds. Thereafter, the cells were harvested for assays.

### ELISA Assays

To determine the extent to which various compounds could suppress extracellular matrix generation, we assessed the generation of fibronectin by the TGF-β1 stimulated fibroblasts after overnight culture with the various test agents. On day one of the ELISA assay, the cells on the 96 well culture plate were washed with PBS and then lysed with 125 μl of high detergent RIPA buffer with protease inhibitors. The cell lysates were transferred to an ELISA plate along with a fibronectin standard curve and protein allowed to bind for 12 hours at 4°C. On day two of the assay, the ELISA plates were washed with PBS-T (PBS containing 1% TWEEN) and then a blocking agent consisting of 3% BSA was applied for one hour. Thereafter, an anti-fibronectin antibody (Sigma-Aldrich, St Louis, MO) derived from rabbit was applied to the plate and incubated at room temperature for two additional hours. The plates were then washed with PBS-T and an anti-IgG rabbit peroxidase (Sigma-Aldrich, St Louis, MO) was applied and incubated at room temperature for one hour. The plates were washed again five times with PBS-T and developed using a TMB substrate solution (eBioscience, San Diego, CA) for twenty minutes. Finally, a stop solution of 2N sulfuric acid was applied. The plates were then measured on a spectrophotometer reading at a wavelength of 450 nm.

### XTT cell viability assays

To verify that the various test agents were not reducing fibronectin generation by exerting cell toxicity, we further performed XTT cell viability assays in the presence of the test compounds. The cell viability assays were performed after the cell culture conditions as outlined above. The Cell Proliferation Kit II (XTT) was supplied from Roche, Inc. The components of the kit were stored at −20°C and protected from light. The buffers, XTT labeling reagent, and electron coupling reagent, were thawed immediately before use at 37°C and mixed thoroughly in each vial to obtain a clear solution. The XTT labeling mixture was then prepared by mixing 50 μl XTT labeling reagent and 1 μl electron coupling reagent and applying 50 μl of the mixture to the cell culture plate and incubating at 37°C in a humidified atmosphere for four hours. After this time period, the absorbances were then measured at 450-500 nm with a reference wavelength of 650 nm. Drug compounds were considered non-toxic if the cell viability was at 80% or greater ^13^.

### Statistical analysis

Statistical analysis was performed using Prism version 5.0b software (GraphPad, Inc.). Initial ANOVA comparisons were made across the various single drug doses and the control sample of TGF-β1 plus DMSO. Subsequently, two sample students T-tests were performed to determine whether statistically significant differences were present between individual drug concentration effects and control. To analyze for potential synergy between test compounds and nintedanib, half maximal effective doses concentrations were determined and tested alone and in combination. Analysis for synergistic effect between drug combinations in comparison with single drug suppression level was performed using two sample students T-tests. All differences were considered to be statistically significant when P < 0.05.

## RESULTS

### Fibroblast screening reveals unexpected drugs with antifibrotic activity

From the initial screening library of over 1524 FDA approved drugs, a comparison of the suppression of fibroblast fibronectin generation was made using the control of pirfenidone at a concentration of 5 mM as a baseline, which was the concentration of this compound that reliably suppressed fibronectin production by the cultured fibroblasts ^14–16^. After analysis, 62 out of the original drug compounds from the repurposing drug screening library at concentrations of 10 μM were shown to suppress fibronectin expression greater than pirfenidone at a 5 mM concentration and did not result in significant cell toxicity when assessed by XTT cell viability assay. Specifically, drug compounds within the category of NSAIDs, steroids, azole antifungal agents, Vardenafil, and antipyrine were associated with significant suppression of the extracellular matrix protein fibronectin on ELISA assay (Table 1). These observations indicate that drug screening can reveal unexpected compounds with novel antifibrotic activity *in vitro*.

**Table 1:**
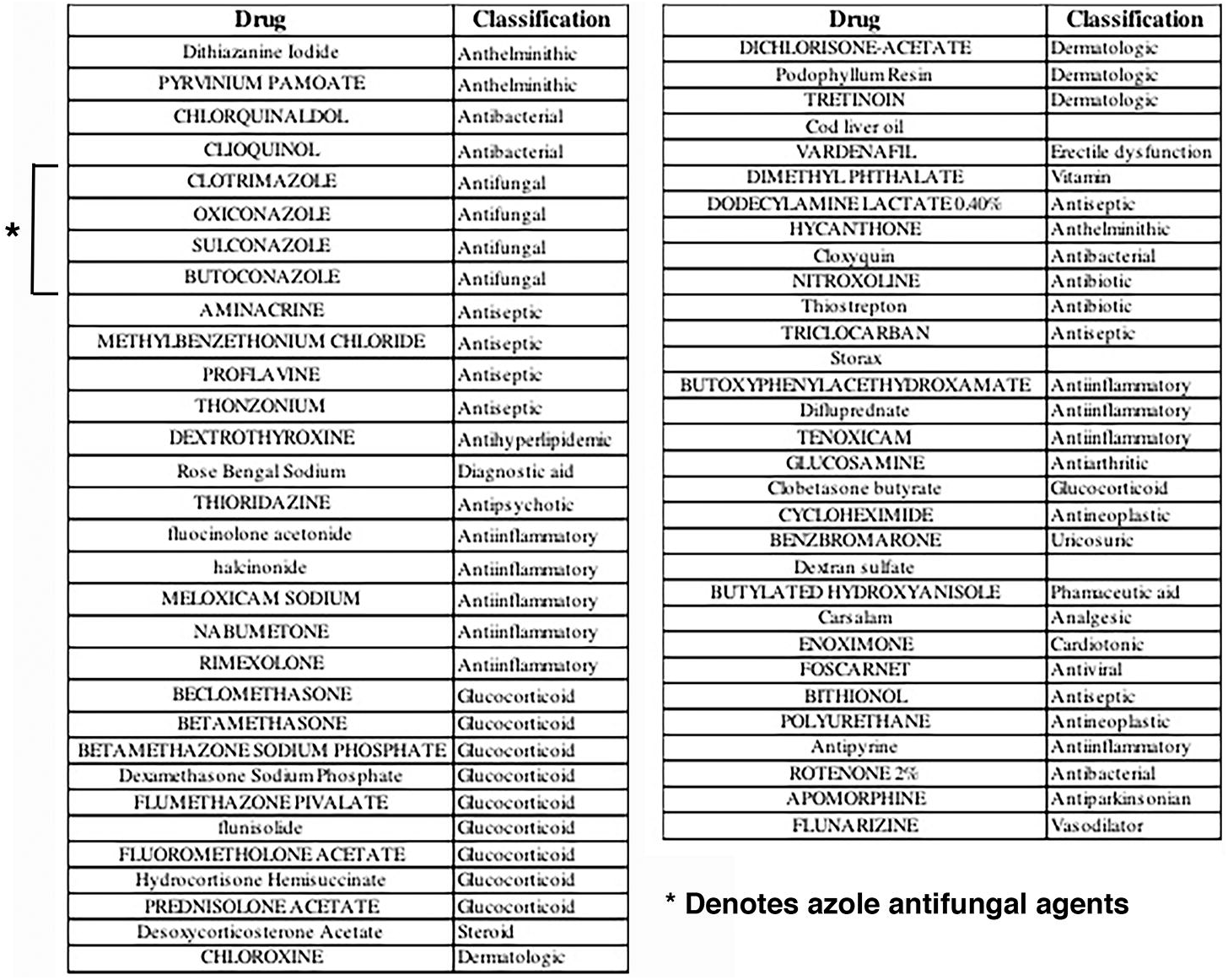
Agents following screening that possessed potential antifibrotic activity

### Certain azole antifungals demonstrate antifibrotic activity on cultured fibroblasts

Of the azole antifungal agents tested, oxiconazole, clotrimazole, and butoconazole showed significant fibronectin suppression (Figure 1). Oxiconazole demonstrated significant fibronectin suppression at 10 μM and 1 μM concentrations (P= 0.0103 and 0.018, respectively). Clotrimazole also demonstrated significant fibronectin suppression at 1 μM and 0.1 μM concentrations (P= 0.023 and 0.0482, respectively). Butoconazole demonstrated significant fibronectin suppression at 10 μM and 1 μM concentrations (P= 0.0282 and 0.0221, respectively). Taken together, these data support that some azole antifungal agents, also have previously unexpected antifibrotic activity suppressing extracellular matrix protein generation by cultured fibroblasts.

**Figure 1:**
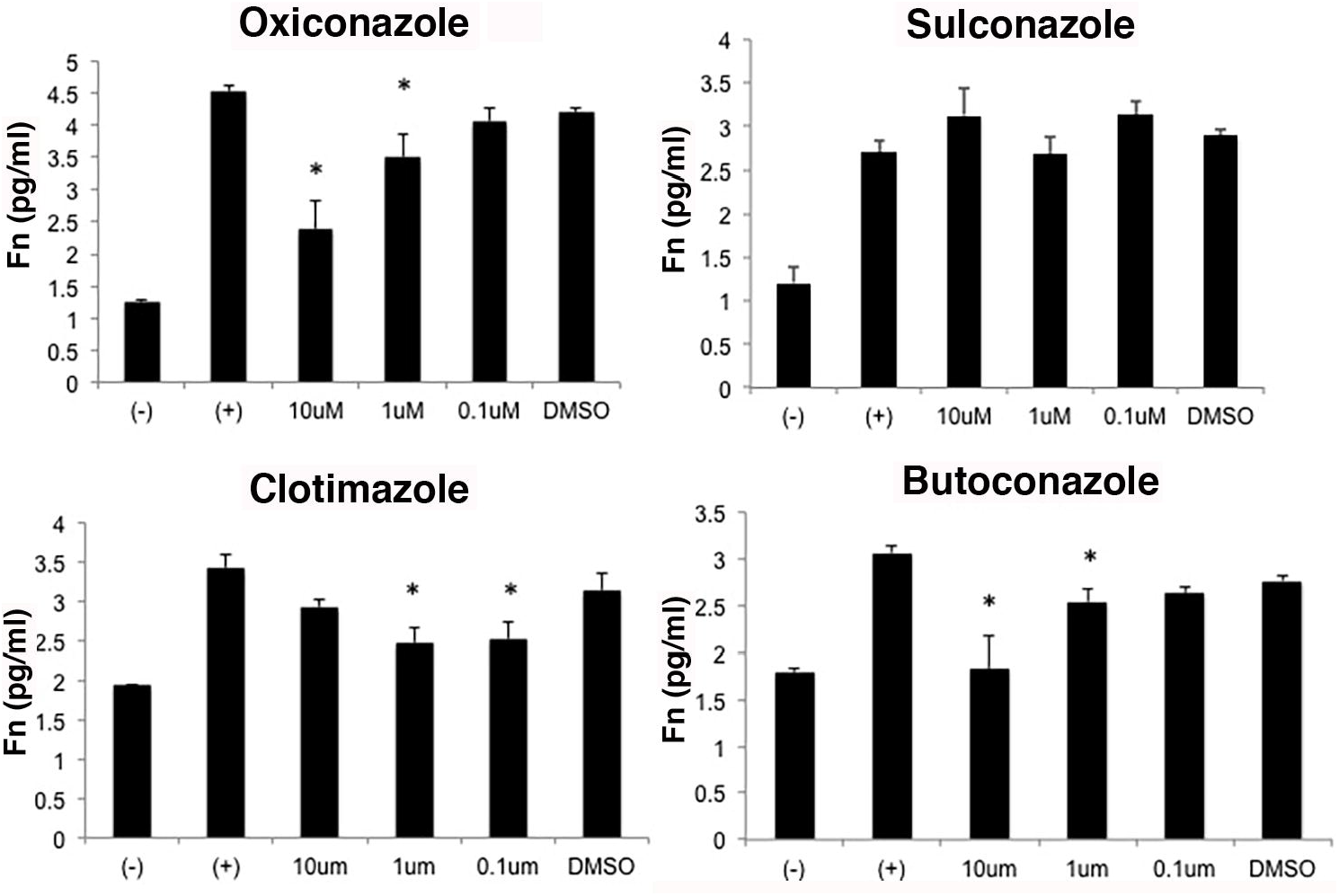
Certain azole antifungals suppress extracellular matrix fibronectin production by TGF-β1 stimulated fibroblasts. AKR-2B fibroblasts were cultured overnight in the presence of the various agents at the indicated concentrations. Similar concentration of DMSO with TGF-β1 served as controls. The following days the cells were lysed and fibronectin (Fn) content assessed by ELISA. (*Denotes P<0.05 compared with controls.)

### Combinations of oxiconazole and pirfenidone demonstrate further suppression of fibroblast fibronectin generation compared to pirfenidone alone

To initially investigate potential drug combinations, we next tested oxiconazole as a prototypic azole in combination with pirfenidone for potential additional anti-fibrotic effect on extracellular matrix fibronectin generation by the cultured fibroblasts (Figure 2). The combination of oxiconazole at a 10 μM concentration with pirfenidone at a 5 mM concentration demonstrated additional suppression of fibronectin when compared to 5 mM pirfenidone alone (P=0.031). The combination of oxiconazole at a 5 μM concentration with pirfenidone at a 5 mM concentration also demonstrated additional suppression of fibronectin when compared to 5 mM pirfenidone alone (P= 0.0047). Furthermore, the combination of oxiconazole at a 1 μM concentration with pirfenidone at a 5 mM concentration demonstrated synergistic suppression of fibronectin when compared to 5 mM pirfenidone alone (P= 0.0161). Thus, combinations of oxiconazole and pirfenidone demonstrate further suppression of fibroblast fibronectin generation compared to pirfenidone alone.

**Figure 2:**
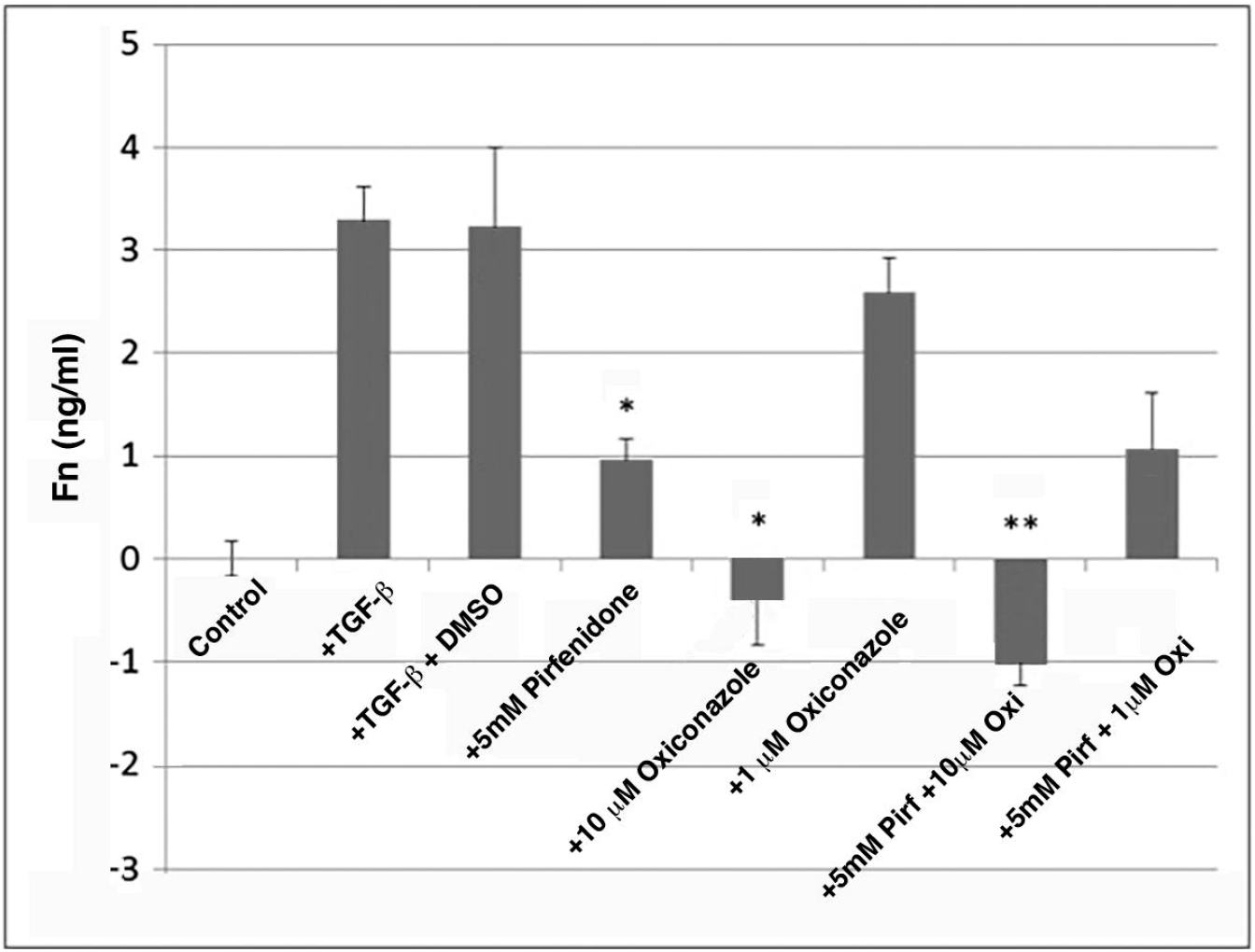
Oxiconazole suppresses extracellular matrix fibronectin production by TGF-β1 stimulated fibroblasts alone and in combination with pirfenidone. AKR-2B fibroblasts were cultured overnight in the presence of the various agents at the indicated concentrations. Similar concentration of DMSO with TGF-β1 served as controls. The following days the cells were lysed and fibronectin (Fn) content assessed by ELISA. (*Denotes P<0.05 compared with controls. **Denotes P<0.05 compared to the level of matrix suppression observed with pirfenidone alone.)

### Combinations of oxiconazole demonstrate further suppression of fibroblast fibronectin generation compared to nintedanib alone

We next evaluated whether oxiconazole in combination with nintedanib could give additional potential anti-fibrotic activity on extracellular matrix fibronectin generation by the cultured fibroblasts (Figure 3). The combination of oxiconazole at 1 μM concentration with nintedanib at 5 μM concentration demonstrated additional suppression of fibronectin when compared to 5 μM nintedanib alone (P= 0.0318). Therefore, once again combinations of oxiconazole and nintedanib further demonstrate additional suppression of fibroblast fibronectin generation compared to nintedanib alone.

**Figure 3:**
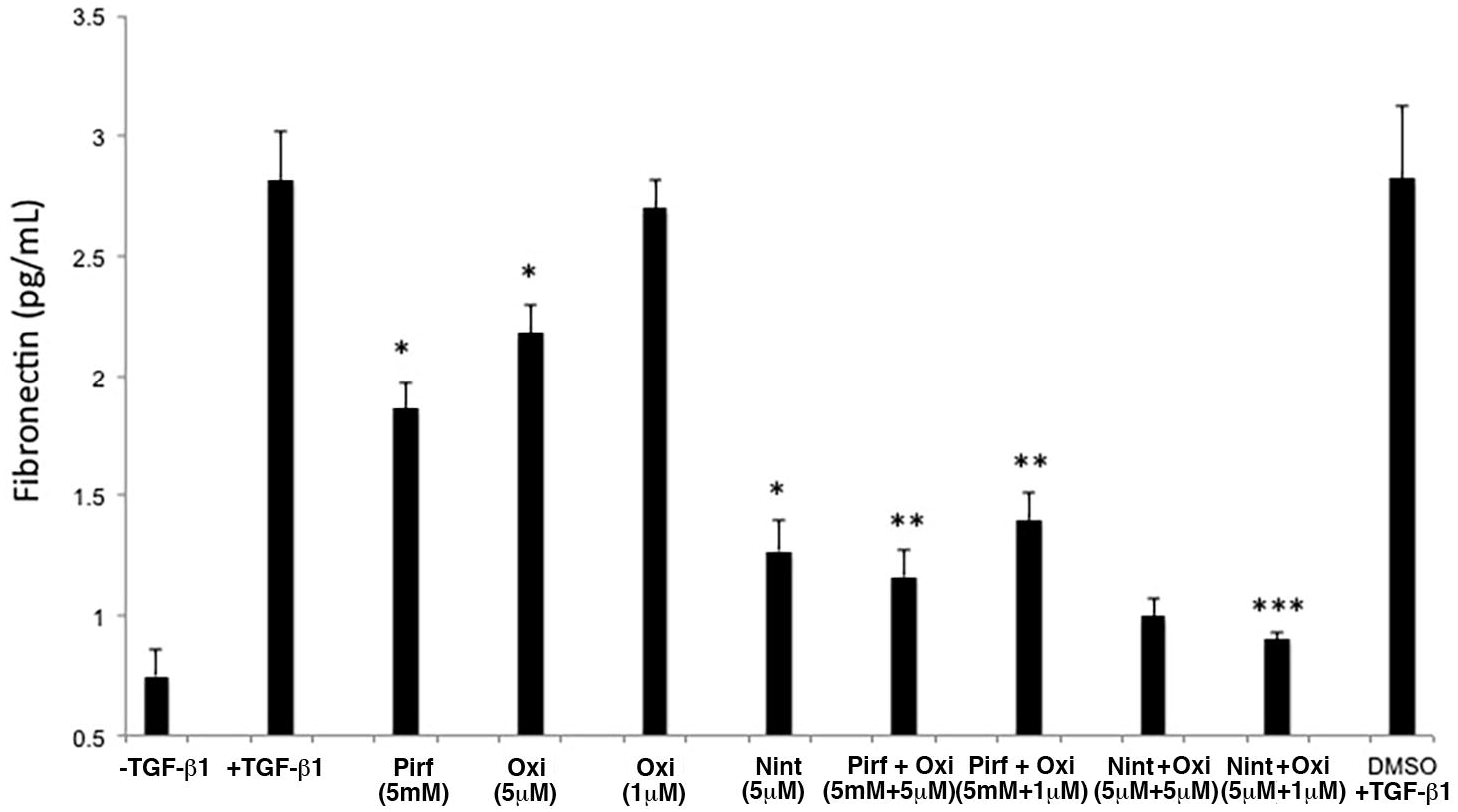
Oxiconazole suppresses extracellular matrix fibronectin production by TGF-β1 stimulated fibroblasts alone and in combination with pirfenidone and nintedanib. AKR-2B fibroblasts were cultured overnight in the presence of the various agents at the indicated concentrations. Similar concentration of DMSO with TGF-β1 served as controls. The following days the cells were lysed and fibronectin (Fn) content assessed by ELISA. (*Denotes P<0.05 compared with controls, indication significant matrix suppression by pirfenidone (Pirf), oxiconazole (Oxi) and nintedanib (Nint). **Denotes P<0.05 compared to the level of matrix suppression observed with pirfenidone alone. *** Denotes P<0.05 compared to the level of matrix suppression observed with nintedanib alone.)

### Oxiconazole demonstrates synergy with nintedanib in suppressing extracellular matrix fibronectin generation by cultured fibroblasts

In light of the extremely high concentrations of pirfenidone required to suppress extracellular matric generation *in vitro*, we further elected to focus on nintedanib for formal synergy testing. Dose response curves were generated that demonstrated synergistic suppression of fibronectin production when oxiconazole and nintedanib are used in combination at dose concentrations of 5 μM and 1 μM (Figure 5; P= 0.0459 and 0.0016, respectively) (Figure 4). Finally, we performed isobole analysis to determine that the combinations of oxiconazole and nintedanib were synergistic rather than simply additive ^11^. To accomplish this the dose effect data of nintedanib and oxiconazole were used and estimated ED50 concentrations for each drug independently determined and a linear isobole model was constructed (Figure 5). Plotting combination of the two drugs (denoted as *) that fell below the line confirmed a synergistic, rather than additive or sub-additive, effect in suppression of fibroblast generation of fibronectin with the combination of oxiconazole and nintedanib (Figure 6).

**Figure 4:**
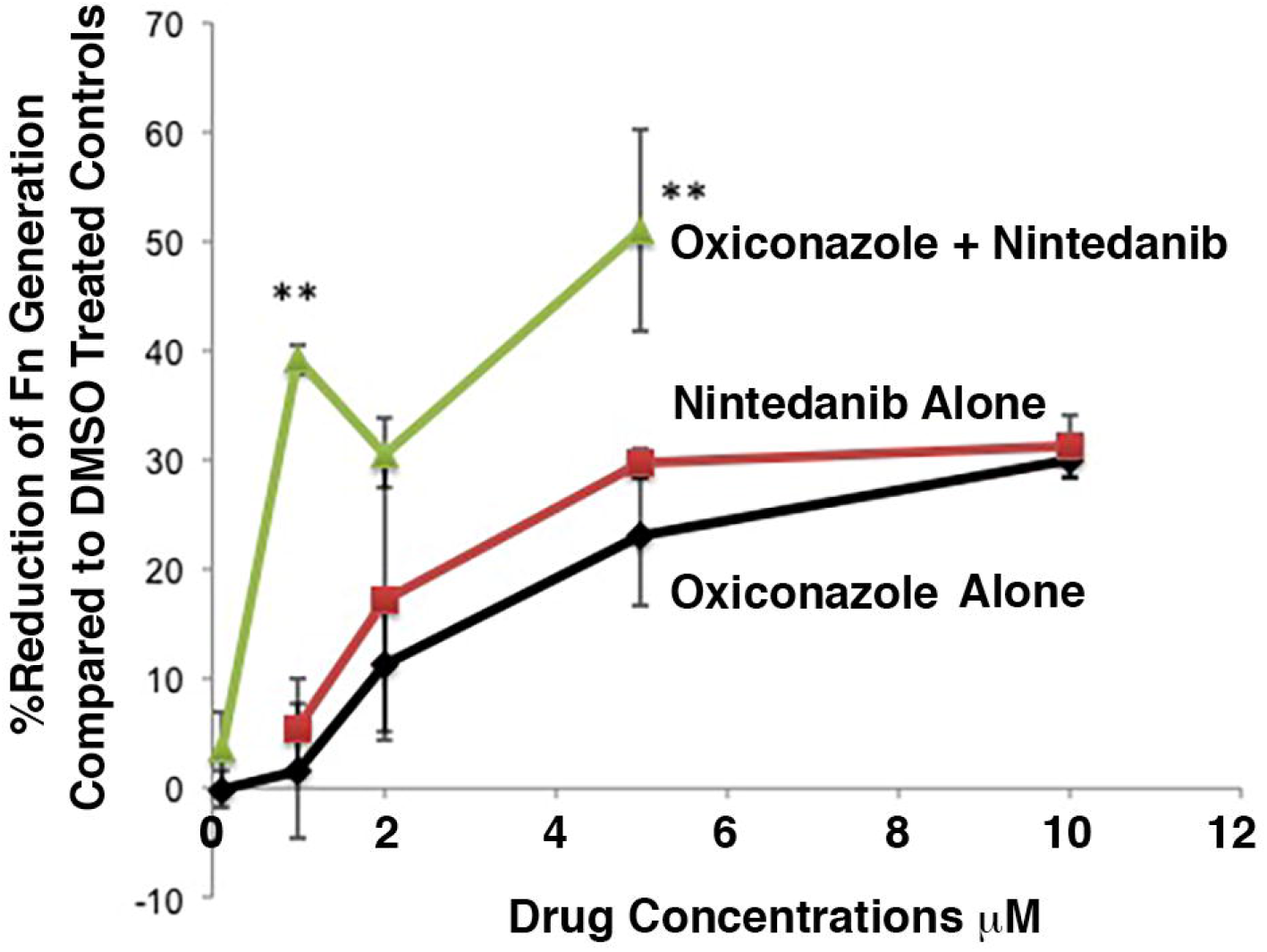
Combinations of oxiconazole with nintedanib suppresses extracellular matrix fibronectin production by TGF-β1 stimulated fibroblasts greater than either agent alone. AKR-2B fibroblasts were cultured overnight in the presence of the various agents at the indicated concentrations. The following days the cells were lysed and fibronectin (Fn) content assessed by ELISA. (**Denotes P<0.05 compared to the level of matrix suppression observed with the combination of oxiconazole with nintedanib compared to either single agent alone.)

**Figure 5:**
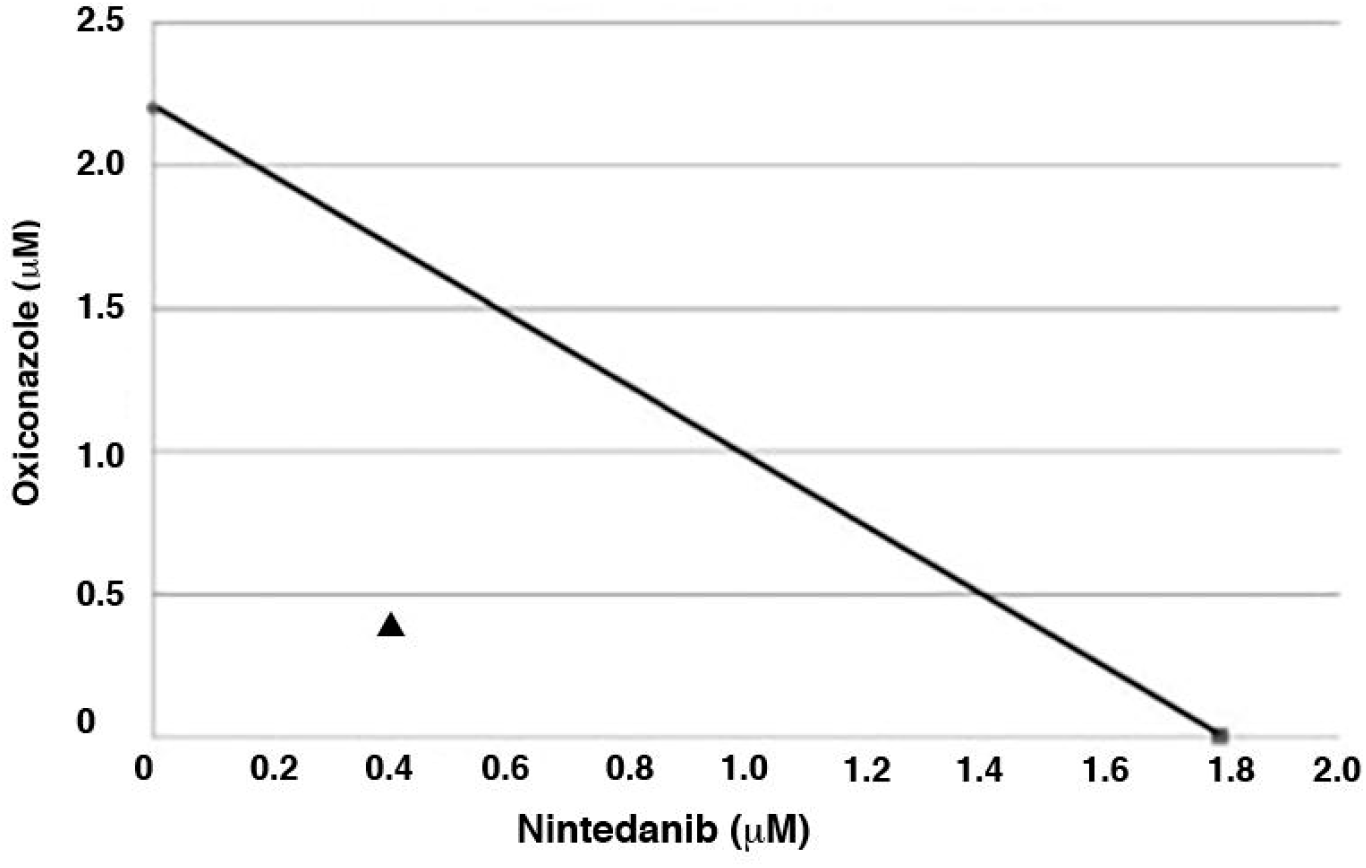
Combinations of oxiconazole and nintedanib were synergistic rather than additive compared to either agent alone. To formally address synergy, the dose effect data of nintedanib and oxiconazole were used and estimated ED50 concentrations for each drug independently determined and a linear isobole model constructed. Plotting combination of the two drugs (denoted as ▲) that fell below the line confirmed a synergistic, rather than additive or sub-additive, effect in suppression of fibroblast generation of fibronectin with the combination of oxiconazole and nintedanib.

## DISCUSSION

This study provides the results of a repurposing drug library screen to identify potential anti-fibrotic agents. The screen revealed a variety of potential drug classes not previously well known for antifibrotic activities. These included the azole antifungal agents, some non-steroidal anti-inflammatory medications, the PDE5 inhibitor vardenafil and other drugs. We further demonstrated that some azole antifungal agents, such as oxiconazole, when used as a test compound could demonstrate synergistic suppression of fibroblast mediated extracellular matrix generation in combination with the currently FDA approved antifibrotic agent nintedanib using an *in vitro* test system.

Initial studies into the use of combination therapy for treatment of IPF have already been attempted. Unfortunately, some combinations have resulted in increased risk of death such as was found for the combination of prednisone, azathioprine and NAC compared to placebo ^17^. Furthermore, the PANORAMA study explored the potential benefit of the addition of acetylcysteine to pirfenidone in patients with IPF. However, this combination therapy also failed to demonstrate improvement in the tolerability profile of pirfenidone and in fact demonstrated a possibly harmful effect with an accelerated rate of decline in forced vital capacity in the combination treatment arm ^18^. Investigations into the combination of pirfenidone and nintedanib is also currently underway and results from the INJOURNEY trial support a manageable safety and tolerability profile of nintedanib with add-on pirfenidone in patients with IPF, similar to those of the individual drugs alone ^19^. This trial also suggested a smaller decline in FVC over the 12-week period in the nintedanib with add-on pirfenidone group. However, the trial was not adequately powered to assess efficacy as an endpoint ^19^. Finally, a recent study demonstrated that the combination of nintedanib with sildenafil, in patients with IPF and a diffusing capacity of 35% or less did not provide a significant benefit as compared with nintedanib alone ^20^.

Currently our understanding of the pathogenesis of disease initiation and progression in IPF remains limited. One hypothesis is that environmental triggers such as microaspiration, tobacco smoke, inhalational exposures, and infections (viral, bacterial, and fungal) all can contribute to the repetitive microinjury of the alveolar epithelium resulting in pulmonary fibrosis ^3,21^. Previously it was believed that the lungs were a sterile environment, but we now know that this is not in fact the case. The lungs have their own microbiome that includes various bacteria, viruses, and fungi ^21^. It has been postulated that alterations to this microbiome of the lung can contribute to exacerbations of and/or disease progression in IPF ^22^. Molyneaux et al. investigated the role of bacterial burden in IPF and identified that patients with IPF have an increased pulmonary bacterial load in bronchoalveolar lavage samples compared to matched controls. They also demonstrated that there was a higher BAL bacterial burden in those patients with IPF with disease progression (defined as a relative decline in FVC of 10% over 6 months or death) compared to those subjects with stable disease ^23^.

Shulgina et al. demonstrated that the addition of twice daily co-trimoxazole compared with standard treatment in a multi-center randomized trial of patients with IPF resulted in a significant reduction in all-cause mortality and reduced the occurrence of respiratory tract infections over a 12 month period ^22^. Yet, it remains difficult to discern whether this effect was the result of the antimicrobial effects of co-trimoxazole or whether the regimen exerted other concurrent anti-inflammatory effects. Also of interest, the PANTHER trial studying the effects of prednisone, azathioprine and NAC therapy actually demonstrated an increased mortality rate for IPF patients treated with immunosuppressive therapies raising the question of increased susceptibility to infections as a contributor to disease progression and mortality ^17^.

The role of fungal infections in the pathogenesis and acute exacerbations of IPF is even less well understood. Previous studies have demonstrated that *Pneumocystis jirovecii* can be associated with acute deterioration in IPF patients and have documented a higher colonization rate in patients with IPF ^24^. Prior postmortem studies of patients with IPF who underwent autopsy have identified fungal infections including bronchopneumonia that were not clinically diagnosed pre-mortem ^24^. While there are potential roles for fungal colonization and infection in the course of IPF, the current study demonstrated previously unrecognized potential antifibrotic activity of certain azole antifungal agents, that occurred *in vitro* in the absence of any fungal organisms or components.

The primary mechanism of action of the azole antifungal agents is known to occur through inhibition of the cytochrome P450-dependent enzyme lanosterol 14-alpha-demethylase. This enzyme is required for conversion of lanosterol to ergosterol and results in damage to the cell membrane of fungi leading to increased permeability and cell lysis/death. In contrast, the potential antifibrotic mechanisms of certain azole antifungal agents remain unknown. Previous studies have indicated that itraconazole has effects on hedgehog signaling ^25^. Hedgehog pathways have also been potentially implicated in IPF ^26^. Interestingly however, itraconazole was not found to possess significant antifibrotic activity in our *in vitro* screening system. Hence, it is unlikely that hedgehog pathways are strongly implicated in our observed antifibrotic activities of other antifungal azoles.

## CONCLUSIONS

Taken together our findings provide an *in vitro* approach for drug screening to identify compounds with potential activity as antifibrotic agents that may be ultimately exploited to treat pulmonary fibrosis. Our studies indicate that certain azole antifungal agents may possess such antifibrotic activity and may have added benefit in addition to standard antifibrotic therapy such as with nintedanib. The antifibrotic mechanisms of these agents remain elusive. But such an *in vitro* approach may be exploited to identify future combination therapies that may be tested in humans with IPF and other fibrotic diseases.

## Author Contributions

CB participated in study design, data collection, and was a major contributor in writing the manuscript. MG participated in data collection. PJ participated in study design and data collection. MB participated in data collection. JC participated in data collection. TK participated in study design, data collection, and writing of the manuscript. AL participated in study design, data collection, data analysis/interpretation, and was a major contributor in writing the manuscript. All authors read and approved the final manuscript.

## Acknowledgements

The authors appreciate many helpful discussions of members of the Mayo Clinic Thoracic Diseases Research Unit during the course of these studies.

## Abbreviations Used

(IPF): Idiopathic pulmonary fibrosis
(FVC): Forced vital capacity
(TGF-β1): Transforming growth factor beta 1
(NSAIDs): Non-steroidal anti-inflammatory drugs
(ELISA): Enzyme linked immunosorbent assay
(NAC): N-Acetyl Cysteine
(FDA): Food and Drug Administration
(ANOVA): Analysis of variance
(FBS): fetal bovine serum
(DMSO): Dimethylsulfoxide
(PBS): Phosphate buffered saline
(RIPA buffer): Radio immunoprecipitation assay buffer
(TMB): Tetramethylbenzidine
(BAL): Bronchoalveolar lavage

